## Supplementary Information for "Opening and closing of a cryptic pocket in VP35 toggles it between two different RNA-binding modes"

### The probability of cryptic pocket opening controls functional tradeoffs in filovirus immune evasion.

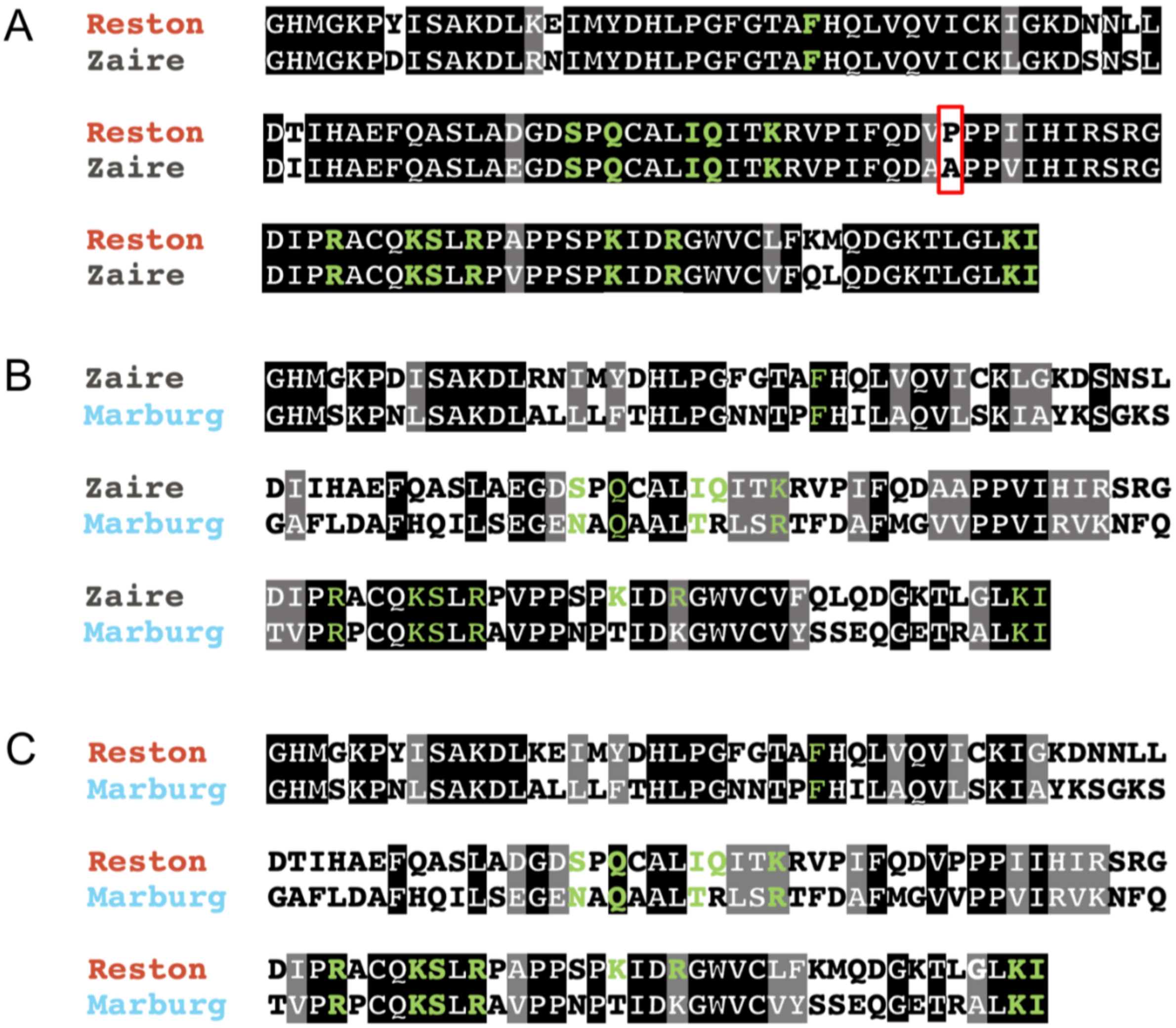


D

| Homolog Pair | Percent Identity | Percent Similarity |
|---|---|---|
| Reston-Zaire | 88.37 | 93.80 |
| Zaire-Marburg | 44.96 | 62.02 |
| Reston-Marburg | 42.64 | 59.69 |

Figure S1: Pairwise sequence comparisons of all homologs used in this study. Residues known to make contacts with dsRNA from structural studies are shown in green. Identical residues are shown with a black background. Similar residues are shown with a grey background. P280 in Reston IID and A291 in Zaire IID are shown in a red box.

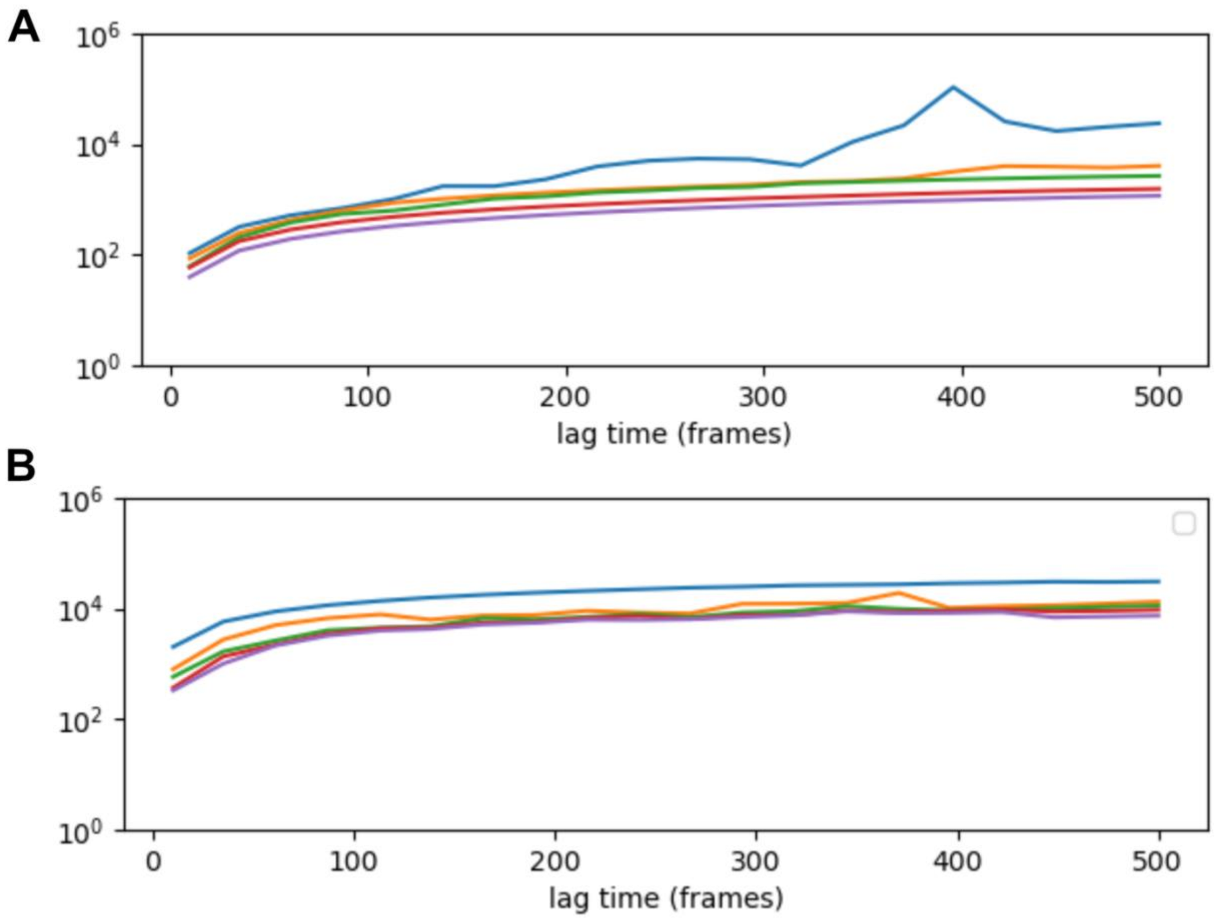


Figure S2: Implied timescales tests for the six slowest eigenvectors of MSMs of Reston IID (A) and Marburg IID (B). Lag times of 6ns were used for both MSMs.

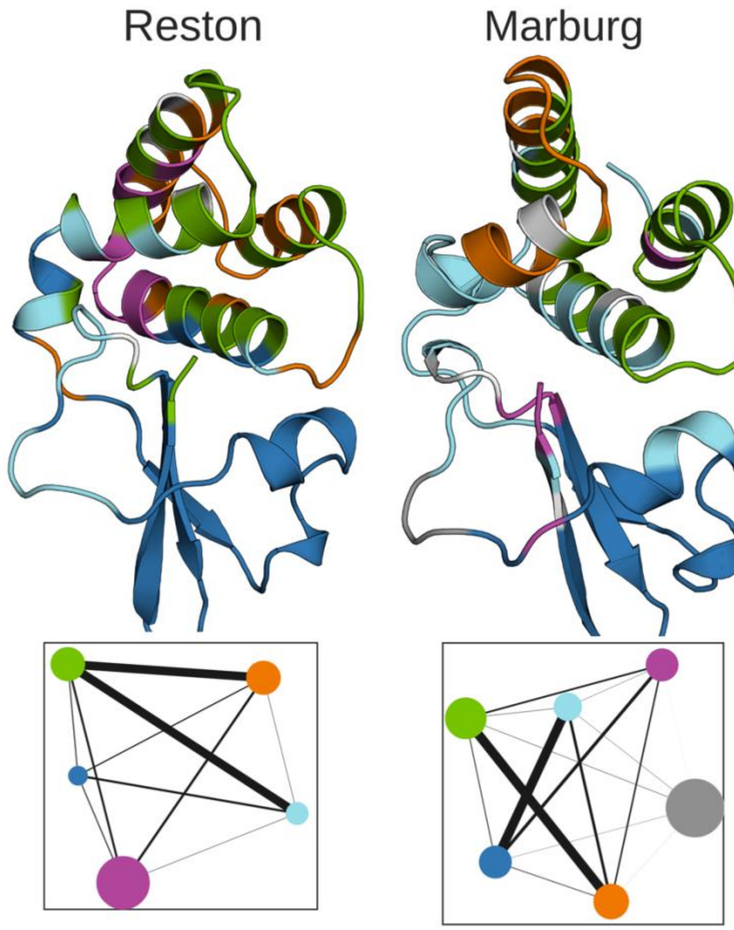


Figure S3: Structure of Reston and Marburg VP35 IIDs with residues in the allosteric network colored according to the CARDs community they belong to. Network representation of the coupling between communities of residues is shown below the corresponding structure colored as in the structures. Node size is proportional to the strength of coupling between residues within the community, and edge widths are proportional to the strength of coupling between the communities.

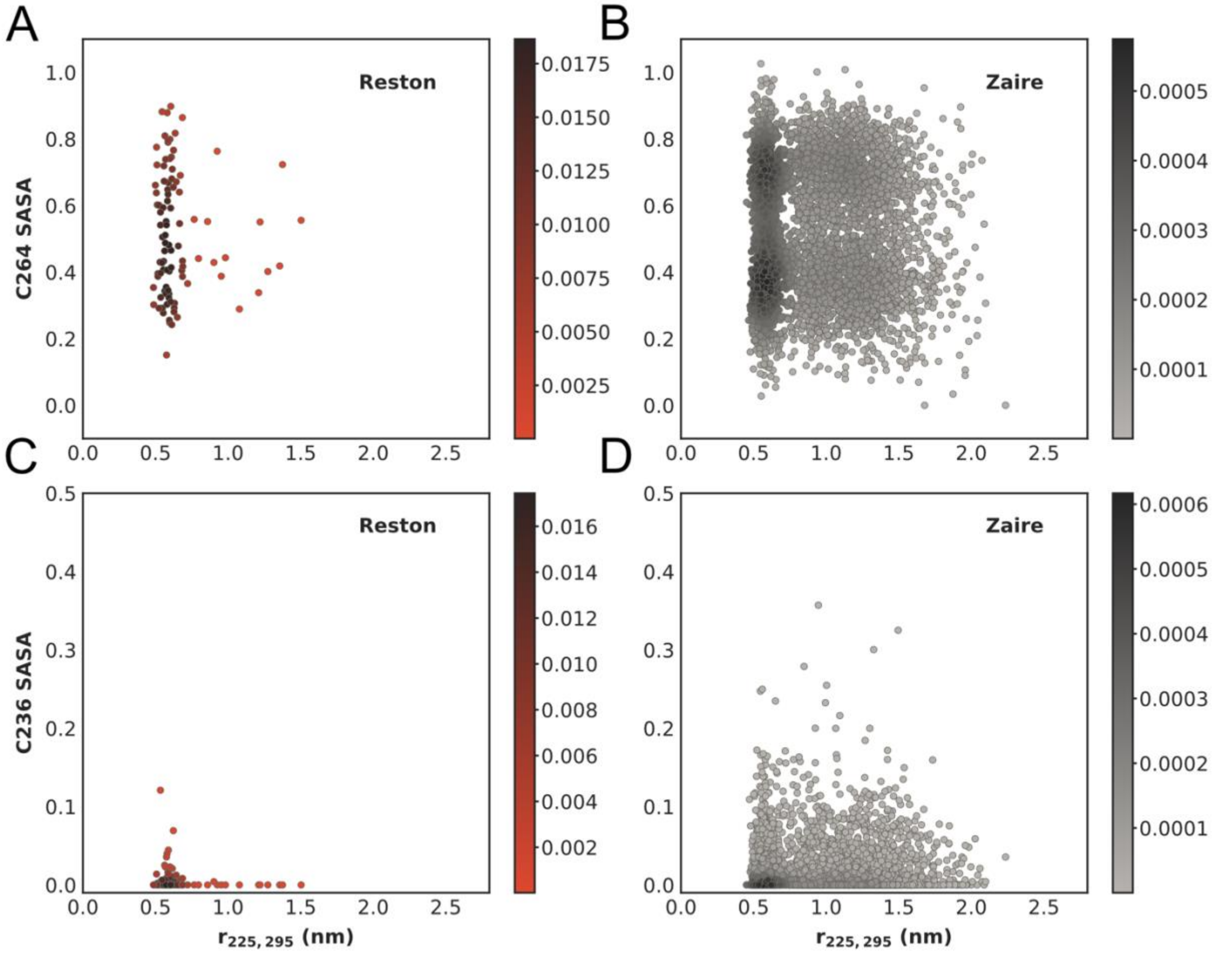


Figure S4: Plots of Solvent Accessible Surface Area (SASA) of C264 of Reston IID(A) and ZAIRE IID (B) and C236 of Reston IID (C) and ZAIRE IID (D) against the distance between residues 225 and 295 calculated from our MSMs.

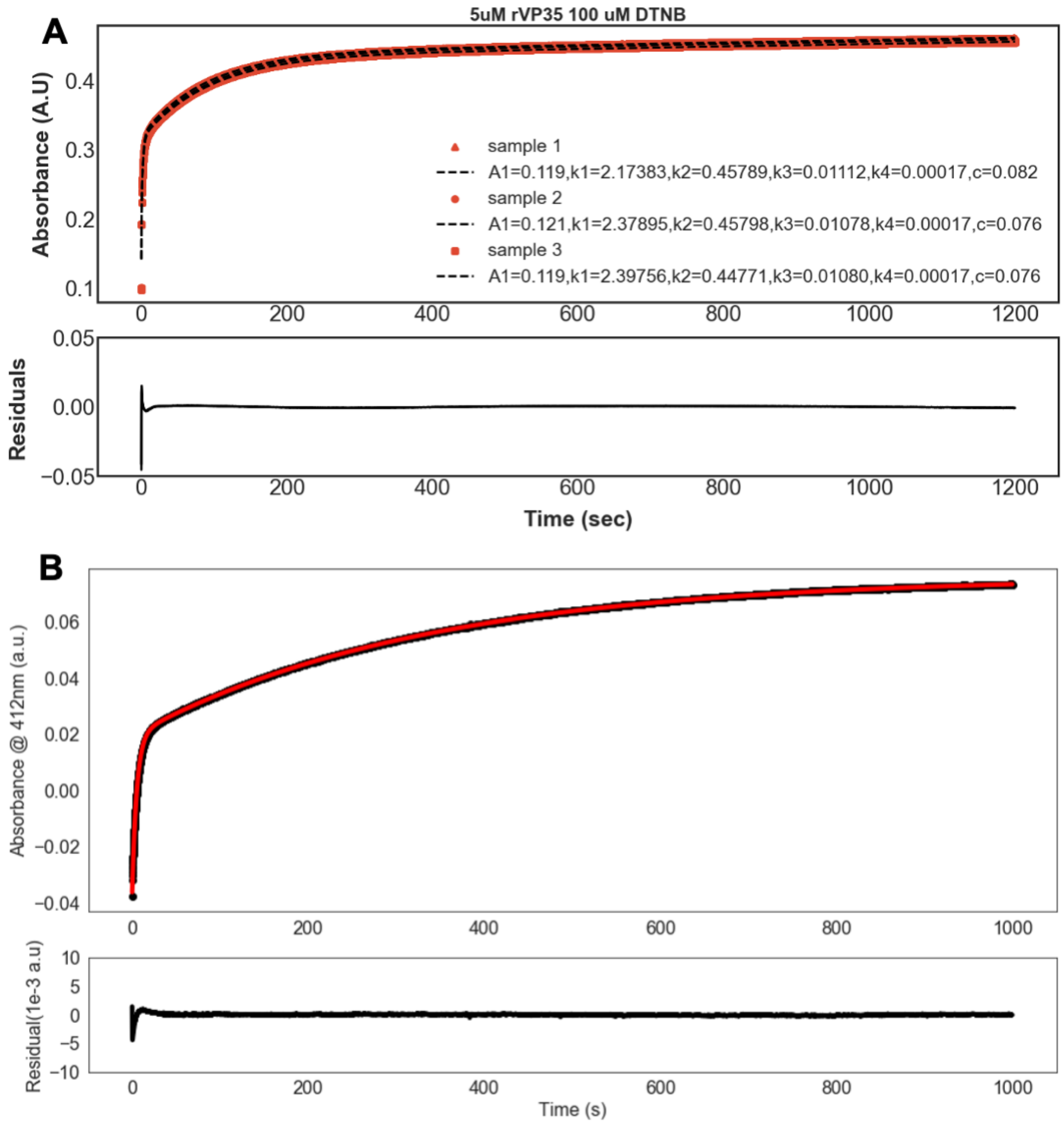


Figure S5 representative time traces from A) three repeats of a thiol labeling experiment (red) performed on Reston IID at 100 μM DTNB and a quadruple exponential fit (black) and B) one repeat of a thiol labeling experiment (black) performed on MARV IID at 100 μM DTNB and a double exponential fit (red). The data are background subtracted (the average absorbance from three runs with DTNB but no protein were subtracted) to account for spontaneous hydrolysis of DTNB

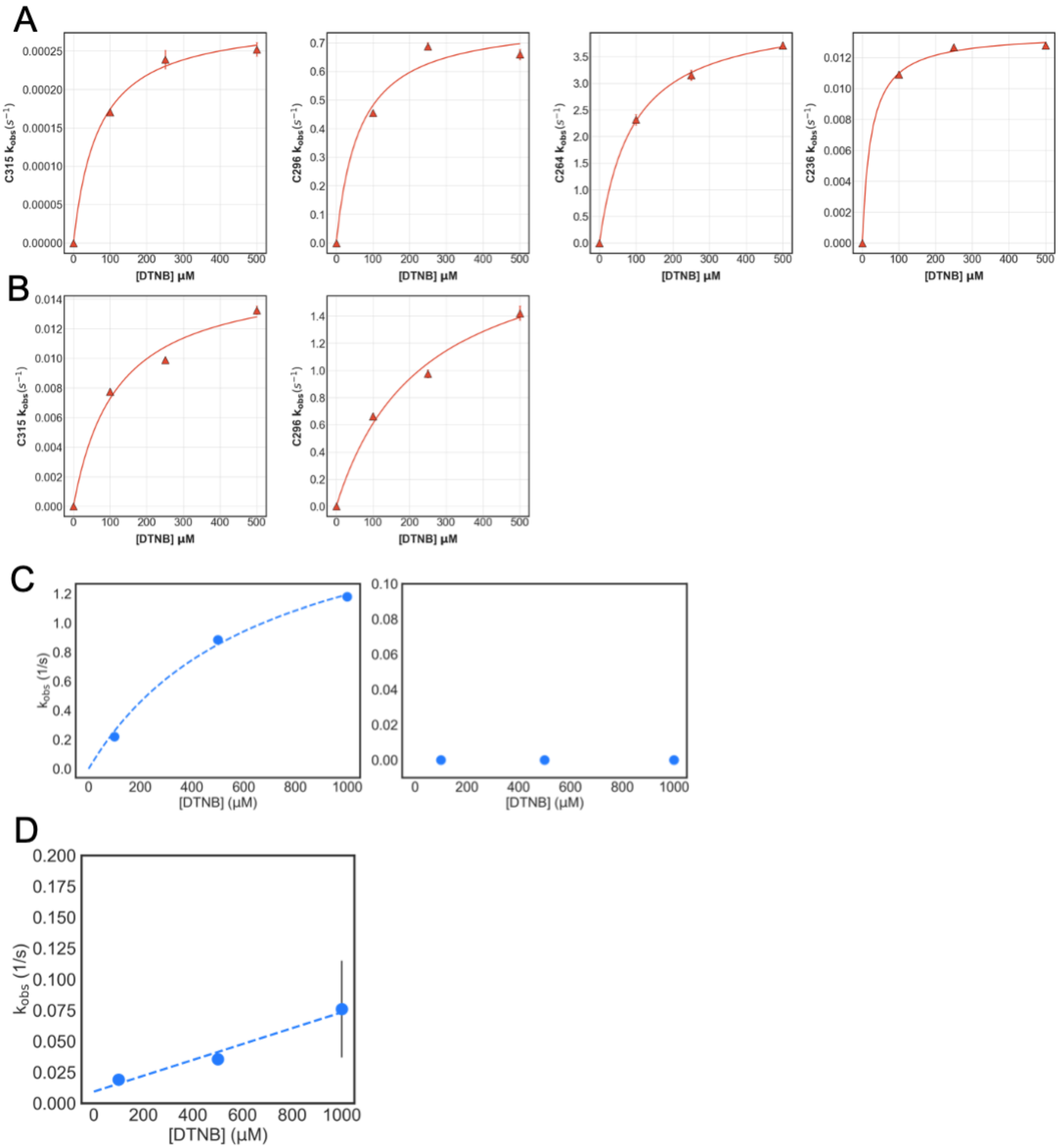


Figure S6 Kobs vs [DTNB] plots from thiol labeling of Reston IID WT (A), Reston IID C236S/C264S (B), MARV IID WT (C) and MARV IID C296S

| | C296 | | |
|---|---|---|---|
| **Homolog** | $k_{open}$ (s-1) | $k_{close}$ (s-1) | $k_{int}$ ($\mu M^{-1} s^{-1}$) |
| **Reston WT** | 0.7652002 ±0.0000004 | 9.7±0.1 | 0.177 ± 0.003 |
| **Reston P280A** | 2.45 ± 0.07 | 1.19 ± 0.01 | 0.047 ± 0.005 |
| **Zaire WT** | 3.17 ± 0.07 | 7.66 ± 0.03 | 0.0136 ± 0.0003 |
| **Zaire A291P** | 0.62 ± 0.04 | 54.63 ± 0.05 | 0.081 ± 0.007 |
| **Marburg WT** | 2.5 ± 0.2 | 0.48 ± 0.005 | 0.0027 ± 0.0002 |

Supplementary Table 1: Estimates for opening ($k_{open}$), closing ($k_{close}$) and intrinsic labeling ($k_{int}$) rates obtained from fits of observed labeling rates as a function DTNB concentration to the Linderstrøm-Lang model.

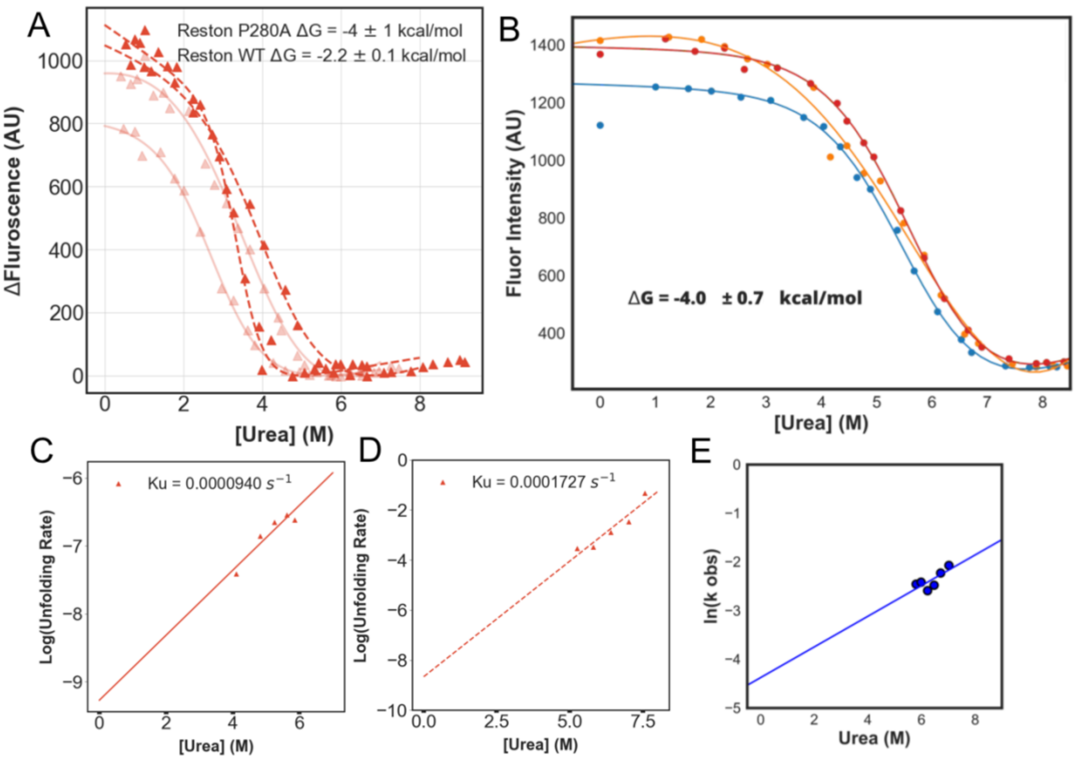


Figure S7: Stability of the homologs obtained from a two state fit to urea denaturation observed via intrinsic tryptophan fluorescence A) Reston WT (light red triangles, solid lines), Reston P280A (dark red triangles, dashed lines), B) Marburg WT. Unfolding rates at 0M urea estimated by measuring unfolding rates at higher urea concentrations for C) Reston WT, D) Reston P280A, and E) Marburg WT. The observed rate for the unfolded fraction is calculated using the Linderstrøm-Lang model using the unfolding rate and the stability of each protein (see Materials and Methods).

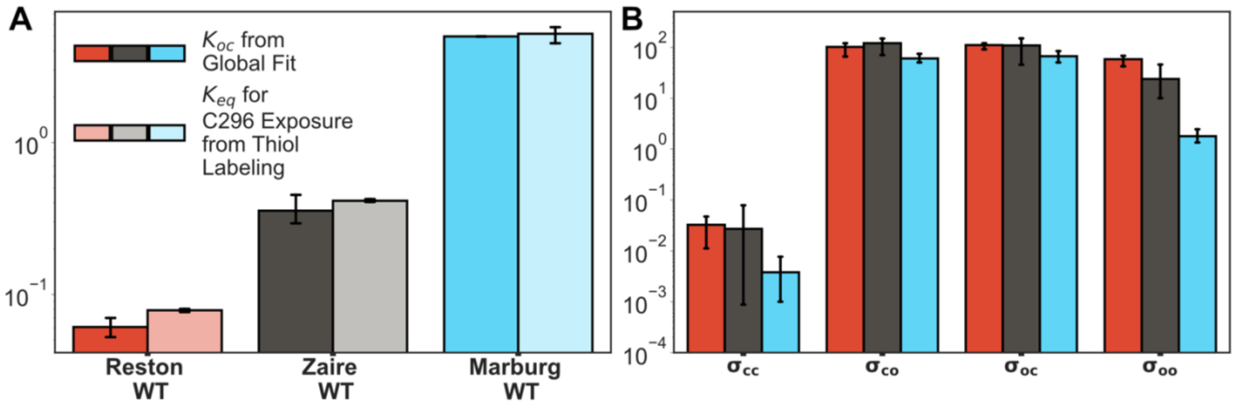


Figure S8: (A) Comparison of $K_{oc}$ obtained from the global fits to RNA-binding data with $K_{eq}$ for C296 (C307 in Zaire IID) exposure obtained from DTNB labeling experiments shown in Fig 3D. (F) Comparison of cooperativity between backbone binding modes obtained from the global fits. Subscript o stands for open state and c stands for closed state. For example, $\sigma_{oc}$ is the cooperativity between the open and the closed states binding in that order along the backbone.

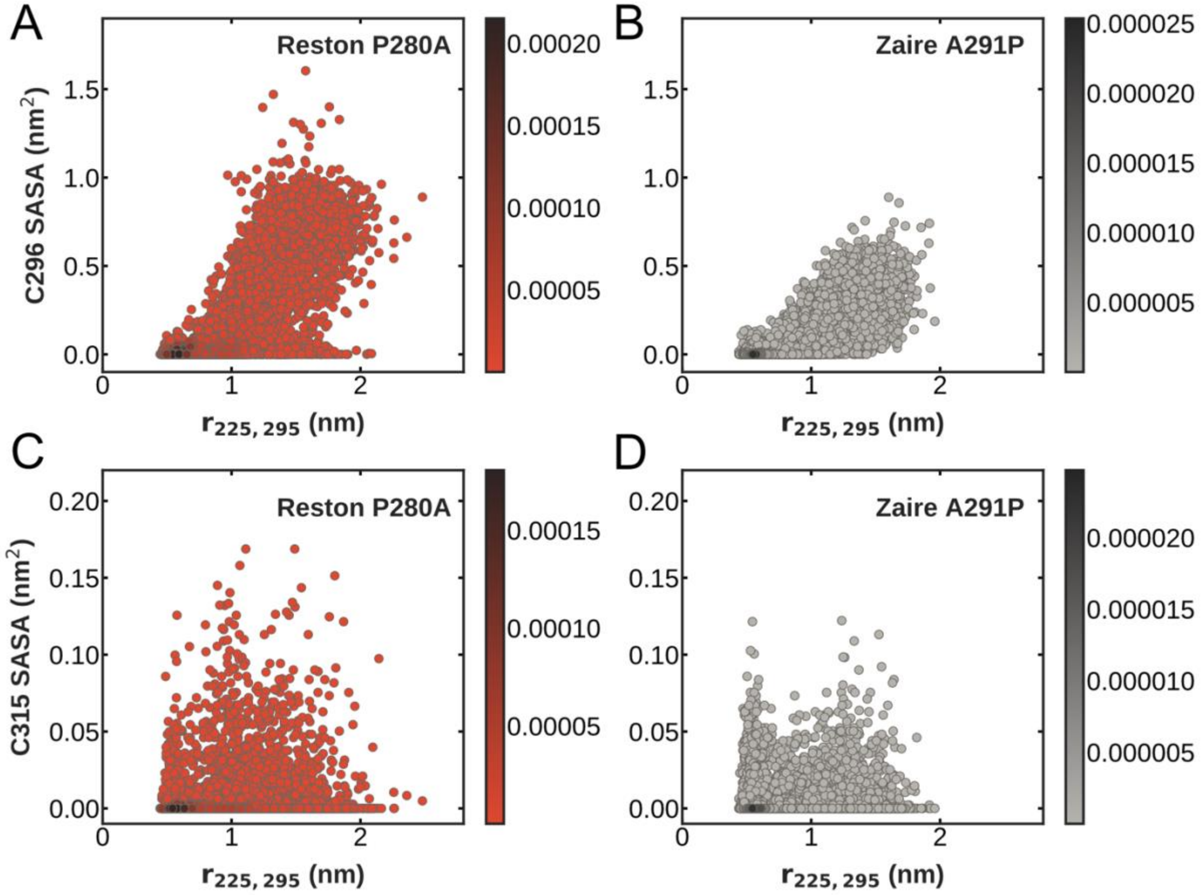


Figure S9: A) Probability distribution of the distance between residues 236 and 306 obtained from MSMs of FAST pockets simulations of Zaire IID WT (solid black) and Zaire IID A291P (dashed black). B) C307 SASA as a function of distance between residues 236 and 306 in Zaire IID A291P C) Observed labeling rates of C307 for Zaire IID WT (transparent solid black) and Zaire IID A291P (dark dashed black). E) C307 SASA as a function of distance between residues 236 and 306 in Zaire IID A291P. F) Observed labeling rates of C326 for Zaire IID WT (transparent solid black) and Zaire IID A291P (dark dashed black)

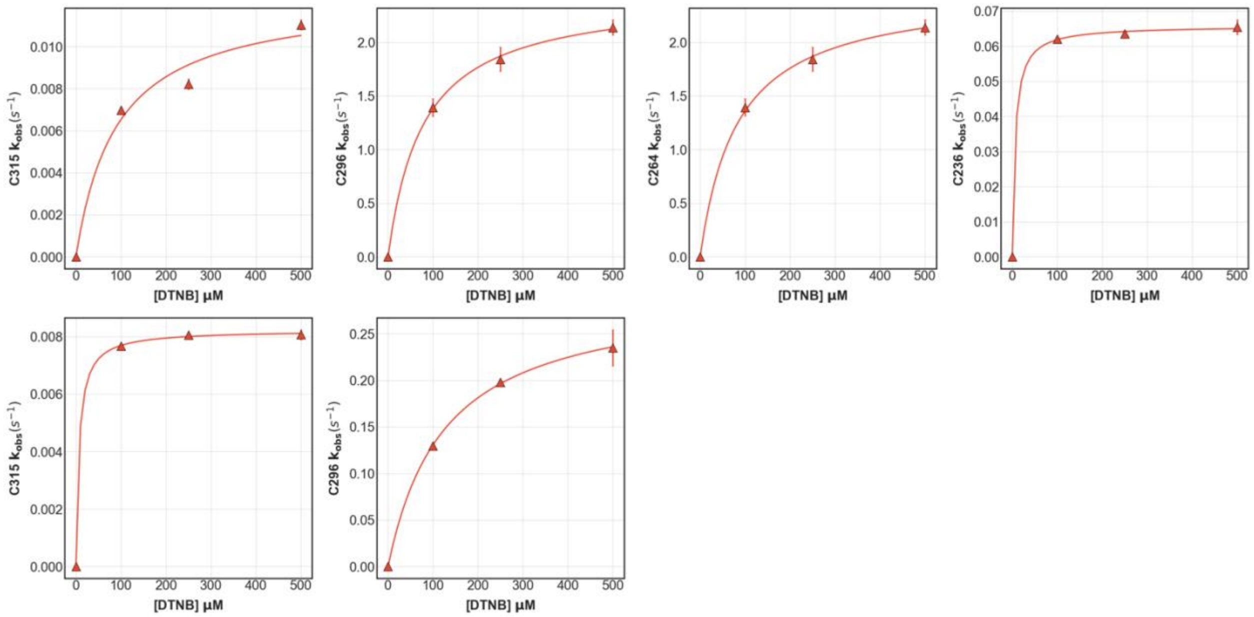


Figure S10 Kobs vs [DTNB] plots from thiol labeling of Reston IID P280A (A), Reston IID P280A/C236S/C264S

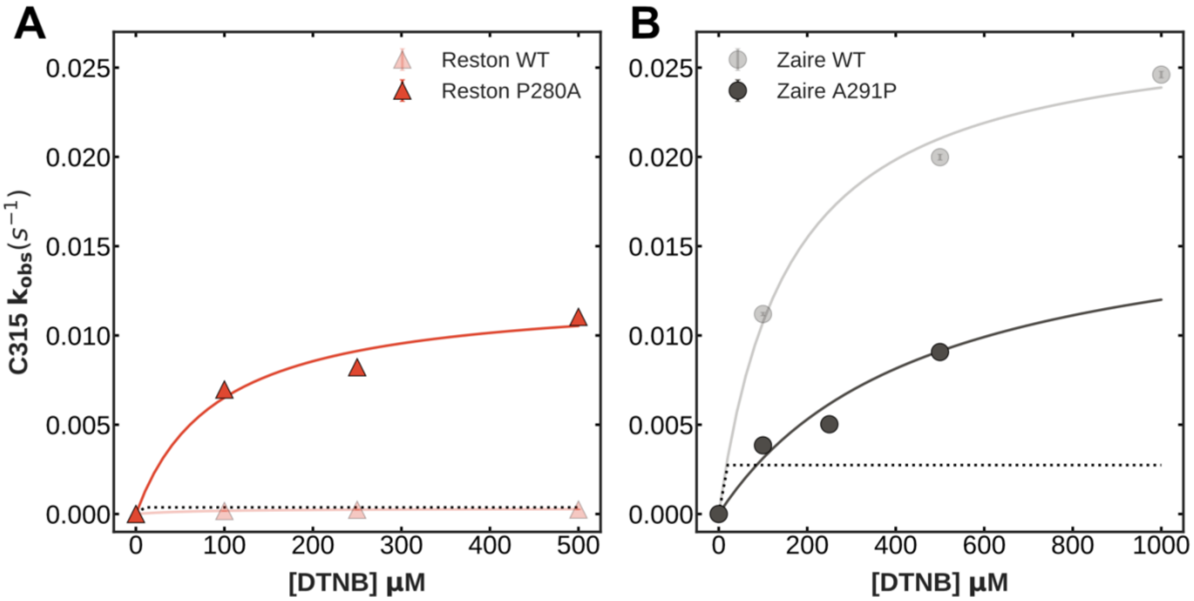


Figure S11: Kobs vs [DTNB] plots for C315 in Reston IID P280A (A) and Zaire IID A291P (B). Both cysteines label faster than expected from the unfolded fraction (shown in dotted lines)

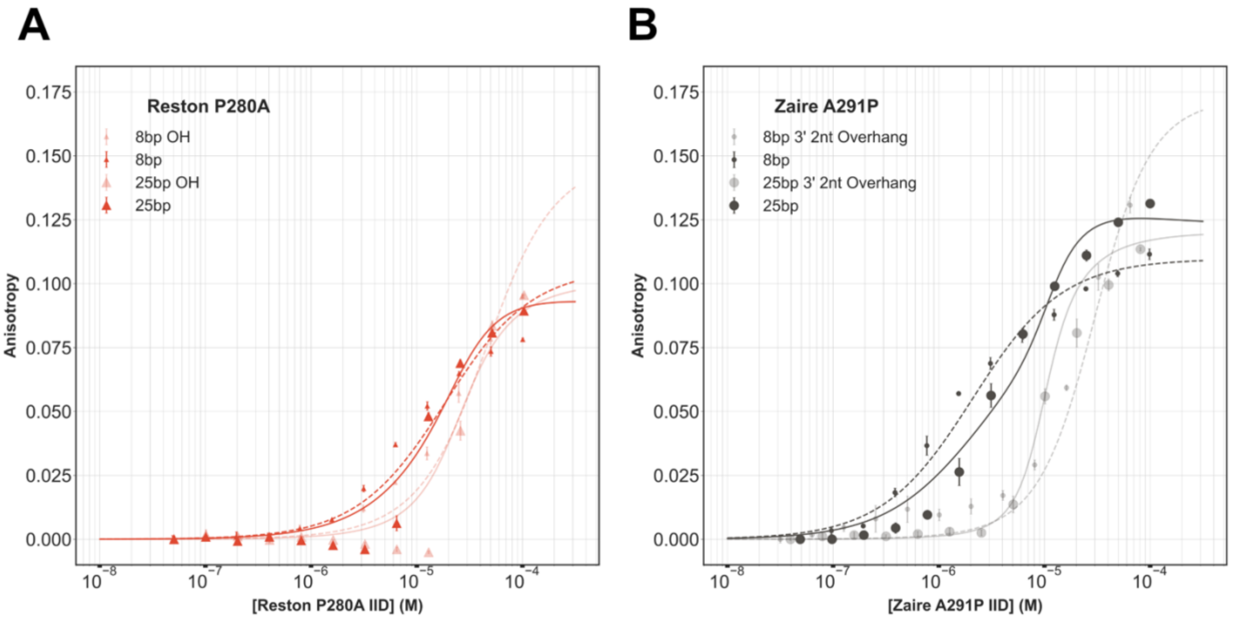


Figure S12: Global fits to the binding model of A) Reston P280A IID and B) Zaire A291P IID.

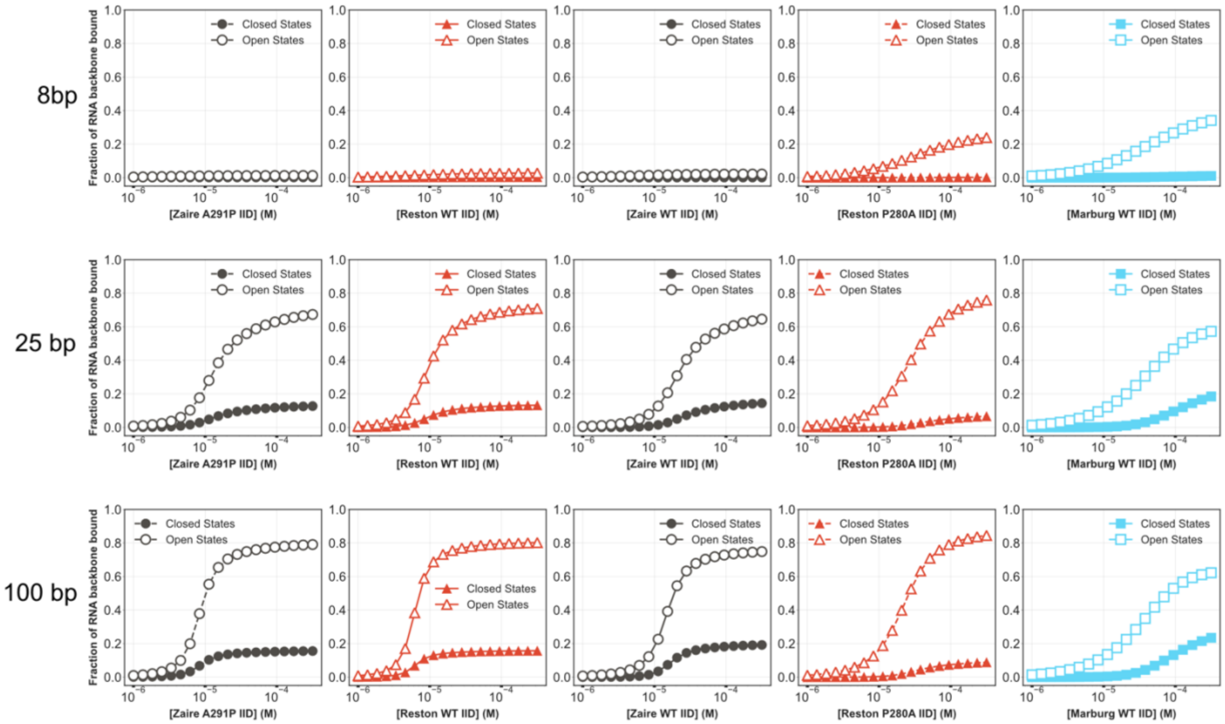


Figure S13: Fraction of the RNA backbone covered by the open and closed states of the various homologs and mutants (in increasing probability of pocket opening from left to right) for 8bp, 25bp and 100bp blunt ended RNA calculated from the binding parameters obtained from the global fits.

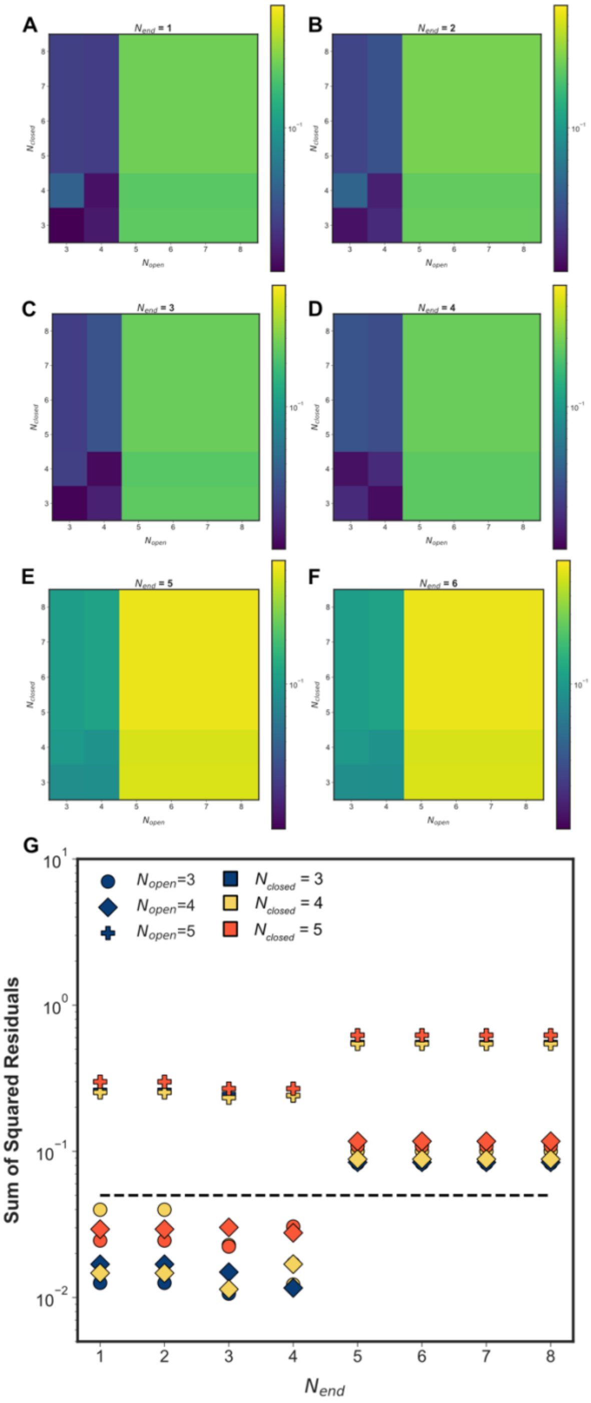


Figure S14: Sum of squared residuals for fits of the RNA binding of all five variants of IID used in this study for various combinations of binding site sizes for the three binding modes. **A-F** show all values of sum of squared residuals (colorbar) for every combination of binding site sizes tested. Horizontal axis in each panel is the binding site size of the open state and vertical axis is the binding site size of the closed state. Each panel shows the matrix for a single end binding site size ranging from 1-6 (A-F) respectively. **G)** Data from A-F plotted to obtain the condition where sum of squared residuals is minimum. End binding site size is plotted on the horizontal axis. Circle, diamond and plus markers indicate backbone binding site size of the open state being 3,4,5 nucleotides respectively. Blue, yellow and orange markers indicate backbone binding site size of the closed state being 3,4,5 nucleotides respectively.

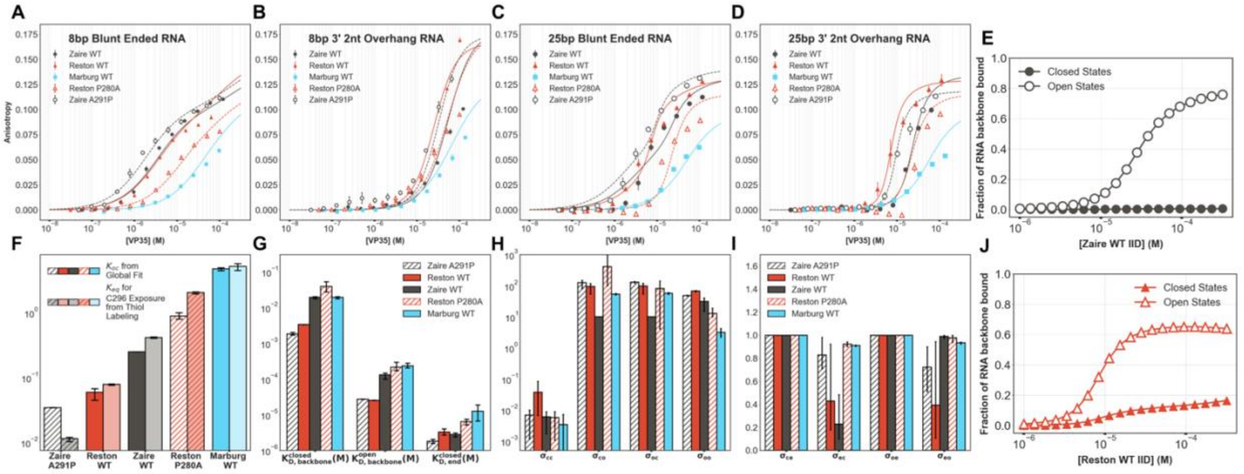


Figure S15: Fits and resulting parameters with the backbone binding site size of the closed state of 3 nucleotides, backbone binding site size of the open state of 4 nucleotides and the end binding site size as 1 nucleotide.

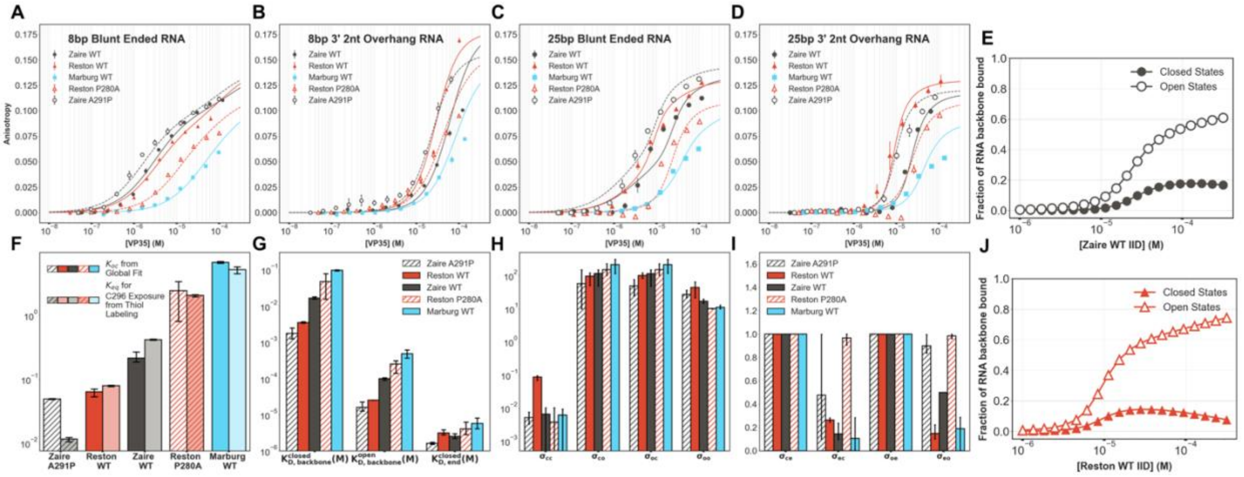


Figure S16: Fits and resulting parameters with the backbone binding site size of the closed state of 4 nucleotides, backbone binding site size of the open state of 3 nucleotides and the end binding site size as 1 nucleotide.

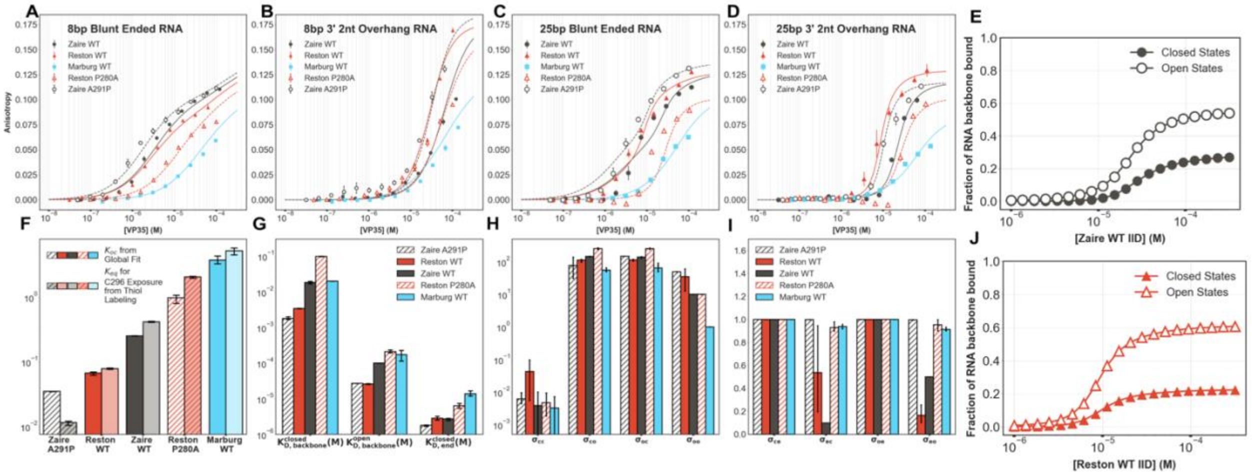


Figure S17: Fits and resulting parameters with the backbone binding site size of the closed state of 3 nucleotides, backbone binding site size of the open state of 3 nucleotides and the end binding site size as 1 nucleotide.

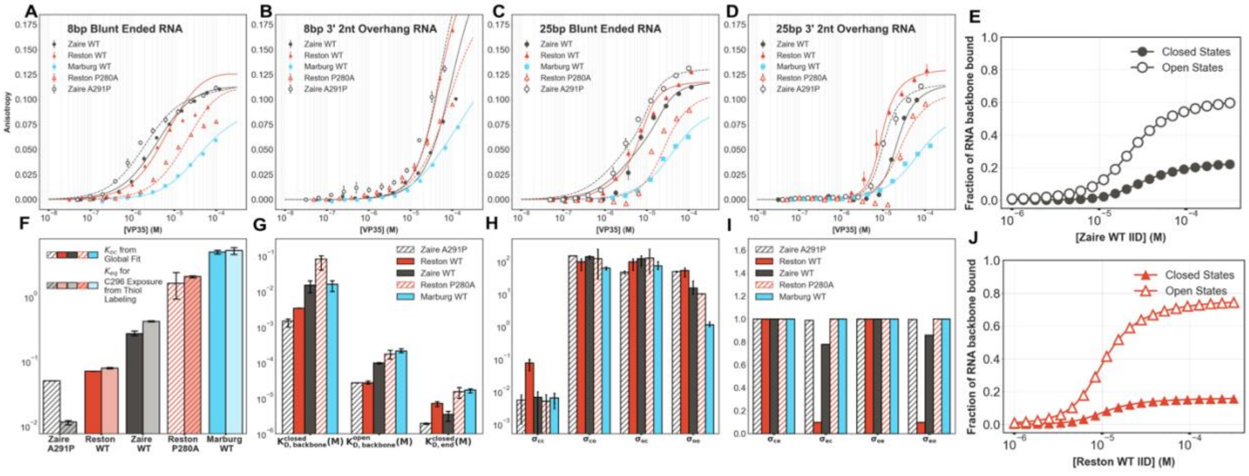


Figure S18: Fits and resulting parameters with the backbone binding site size of the closed state of 4 nucleotides, backbone binding site size of the open state of 4 nucleotides and the end binding site size as 4 nucleotides.

**Interaction** | **Statistical Weight**

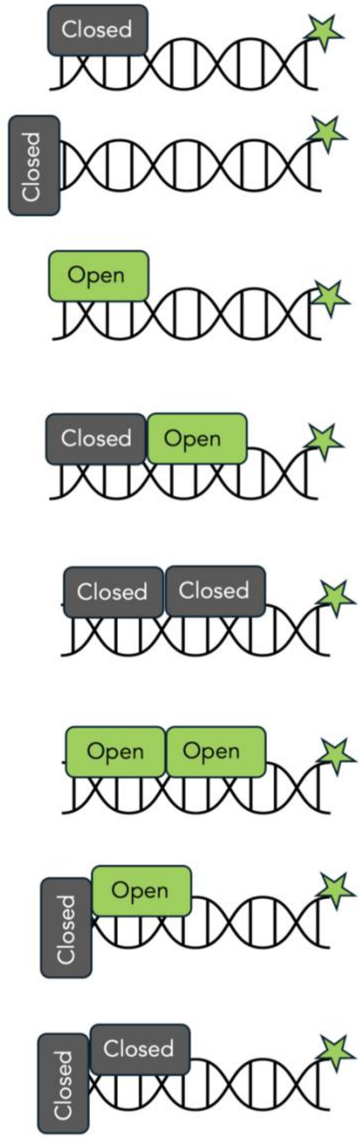


$$\frac{K_A^{closed}[IID]_{total}}{(1+Koc)} \quad (1)$$

$$\frac{(K_A^{end}-K_A^{closed})[IID]_{total}}{(1+K_{oc})} \quad (2)$$

$$\frac{K_{oc}K_A^{open}[IID]_{total}}{(1+K_{oc})} \quad (3)$$

$$\sigma_{co}(1)(3) \quad (4)$$

$$\sigma_{cc}(1)(1) \quad (5)$$

$$\sigma_{oo}(3)(3) \quad (6)$$

$$\sigma_{eo}(2)(3) \quad (7)$$

$$\sigma_{ec}(2)(1) \quad (8)$$

Figure S19: List of different statistical weights for each base pair in the presence of the open and the closed states.

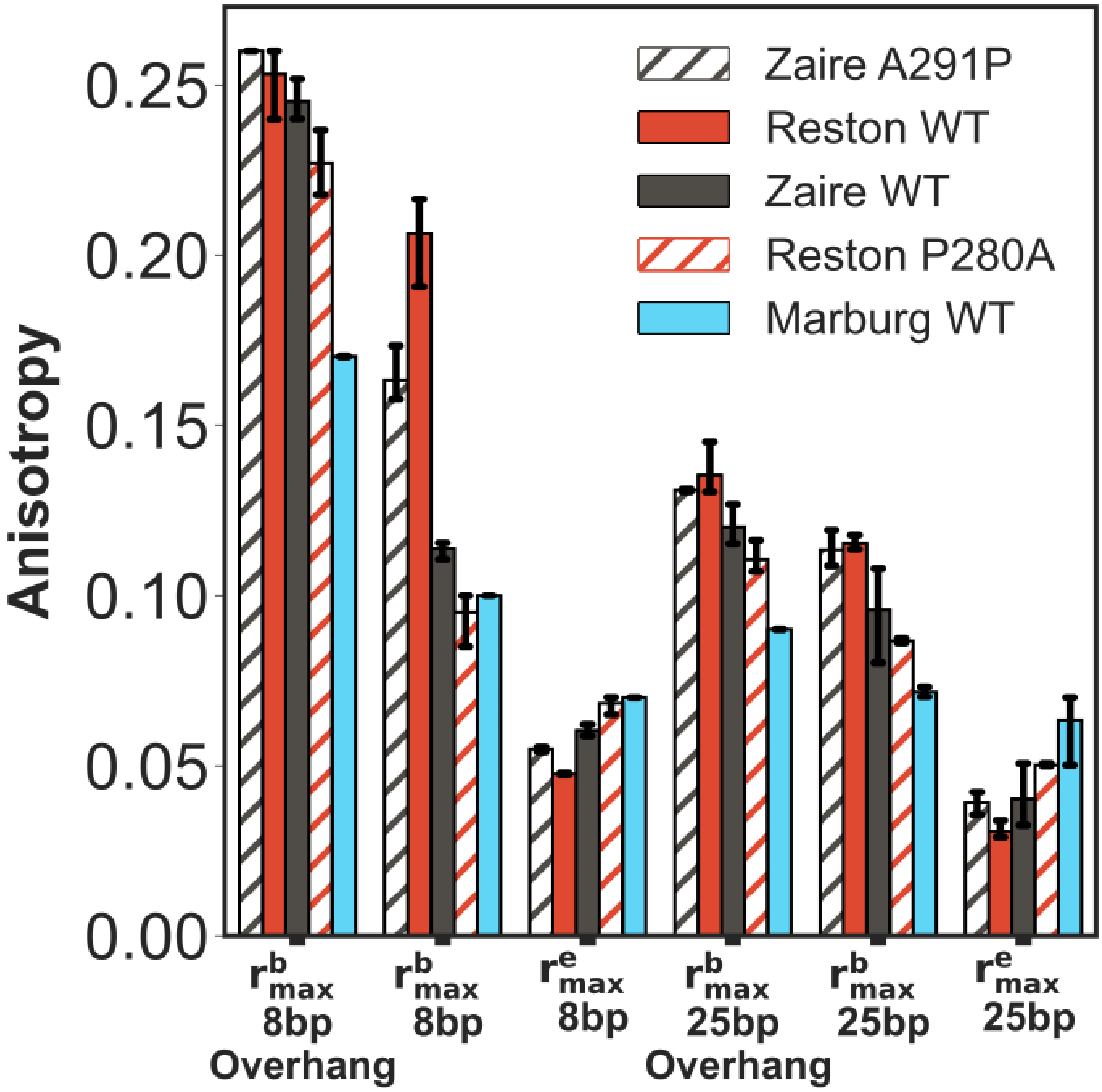


Figure S20 : Maximum anisotropy parameters for the various RNAs obtained the global fits of all five variants of the IID used in this study.
